# Chemogenetic activation of mesoaccumbal Gamma-Aminobutyric Acid projections selectively tunes responses to predictive cues when reward value is abruptly decreased

**DOI:** 10.1101/2020.03.04.977371

**Authors:** Ken T. Wakabayashi, Malte Feja, Martin P.K. Leigh, Ajay N. Baindur, Mauricio Suarez, Paul J. Meyer, Caroline E. Bass

## Abstract

**Background:** Mesolimbic circuits regulate the attribution of motivational significance to incentive cues that predict reward, yet this network also plays a key role in adapting reward-seeking behavior when the contingencies linked to a cue unexpectedly change. Here we asked whether mesoaccumbal gamma-aminobutyric acid (GABA) projections enhance adaptive responding to incentive cues of abruptly altered reward value, and whether these effects were distinct from global activation of all ventral tegmental area (VTA) GABA circuits.

**Methods:** We used a viral targeting system to chemogenetically activate mesoaccumbal GABA projections in male rats during a novel cue-dependent operant Value Shifting (VS) task, in which the volume of a sucrose reward associated with a predictive cue is suddenly altered, from the beginning and throughout the session. We compared the results with global activation of VTA GABA neurons, which will activate local inhibitory circuits and long loop projections.

**Results:** We found that activation of mesoaccumbal GABA projections decreases responding to incentive cues associated with smaller-than-expected rewards. This tuning of behavioral responses was specific to cues associated with smaller-than-expected rewards, but did not impact measures related to consuming the reward. In marked contrast, activating all VTA_(GABA)_ neurons resulted in a uniform decrease in responding to incentive cues irrespective of changes in the size of the reward.

**Conclusions:** Targeted activation of mesoaccumbal GABA neurons facilitate adaptation in reward-seeking behaviors. This suggests that these projections may play a very specific role in associative learning processes.

## INTRODUCTION

Altering the motivational properties of a predictive cue when the value of a reward unexpectedly changes is a critical neurobehavioral component of survival. The mesolimbic system has been heavily implicated in incentive motivational processes, as well as in adapting behavior when reward contingencies change (1-6). While dopamine (DA) neurotransmission from the ventral tegmental area (VTA) to the nucleus accumbens (NAc) is fundamental to both these processes, γ-aminobutyric acid (GABA) neurons are also present in the VTA (7-9). Yet the role of these neurons in regulating reward-seeking behavior is not well understood. While the VTA contains primarily GABA interneurons, there is a subpopulation of VTA_(GABA)_ neurons that project to the NAc (VTA_(GABA)_→NAc, (10-12)). Approximately one third of the total VTA neurons that project to the NAc are GABAergic (13). Optogenetic stimulation of all VTA_(GABA)_ neurons, including interneurons, decreases anticipatory licking in response to an odor predicting a reward during a classical conditioning task (14). Combined with electrophysiology data demonstrating that activation of VTA_(GABA)_ neurons inhibits dopamine neuronal firing during reward expectancy, others have suggested that VTA_(GABA)_ regulation of dopamine neurotransmission contributes to encoding reward prediction error during associative learning processes (14, 15). However, the contribution of VTA_(GABA)_ projections to the NAc in mediating these processes was not evaluated in these studies, and its role remains largely unknown. One study demonstrated that activation of VTA_(GABA)_ terminals in the NAc enhances discrimination between a cue predicting an aversive stimulus (i.e. foot shock) from a non-paired cue via GABAergic inhibition of cholinergic interneurons (CINs) (16). Yet there is scant evidence that VTA_(GABA)_→NAc projections play any role in responding to reward predictive cues. Indeed, we and others have recently reported that directly activating VTA_(GABA)_ terminals in the NAc does not impact sucrose consumption, cue-induced anticipatory licking in mice, or operant responding to reward predictive cues in rats (17, 18).

The primary target of VTA_(GABA)_→NAc projections are tonically active cholinergic interneurons (CINs) (16). CINs “pause” when learning new cue-reward associations (16) and are active when reward-seeking is disadvantageous (e.g. during satiety, (19, 20)). It is therefore possible that VTA_(GABA)_→NAc projections may not have a strong influence on reward-seeking when reward associations are well-established, but are engaged when establishing or adapting to new contingencies. Here we test the hypothesis that increased VTA_(GABA)_→NAc neurotransmission specifically facilitates adapting behavioral responses when the reward value associated with predictive cues abruptly changes. To test this, we chemogenetically activated mesoaccumbal GABA projections selectively during a novel cue-dependent operant task in rats, in which the magnitude of a natural reward (i.e. sucrose) associated with a predictive cue is suddenly altered (Value Shifting, VS). We compared these effects to global activation of all VTA_(GABA)_, which will activate GABA interneurons and projections throughout limbic and cortical brain regions.

## METHODS AND MATERIALS

### Subjects

Long Evans rats (Envigo, Indianapolis, IN) weighing between 290-320 grams were individually housed in 12h/12h light/dark, with lights on at 3AM. Food and water were available *ad libitum*. All procedures complied with the Guide for the Care and Use of Laboratory Animals and were approved by the Institutional Care and Use Committee at the University at Buffalo.

### Adeno-associated virus (AAV)

We use a combinatorial viral method to target specific neuronal subtypes including dopamine (21-23) and GABA neurons (18). Here, we co-infused EF1α-DIO-hM3D-mCherry-AAV2/10 with GAD1-Cre-AAV2/10 into the VTA (n=13, Figure 1A). The expression of the excitatory DREADD hM3D is dependent upon Cre recombinase to reorient the transgene into a translatable position relative to the EF1α promoter. The GAD1-Cre-AAV2/10 construct expresses Cre only in neurons that express glutamate decarboxylase 1. When co-infused they result in hM3D expression in GAD1+ GABAergic neurons (18). It should be noted that the population of dopamine neurons that also make GABA use a biochemical pathway independent of GAD1, and these neurons appear to be GAD1 negative (24). A separate cohort of rats (n=9) was co-infused with GAD1-CRE-AAV2/10 and a CMV-DIO-tdTomato-AAV2/10 to serve as DREADD-free controls. Targeted activation of mesoaccumbal GABA was achieved by microinfusing clozapine-N-oxide (CNO) into the NAc, while global activation of VTA_(GABA)_ occured by systemic administration of CNO (Figure 1A, E). Details of stereotaxic infusions, guide cannula implant surgeries, and CNO treatments are detailed in the Supplemental Materials.

**Figure 1.**
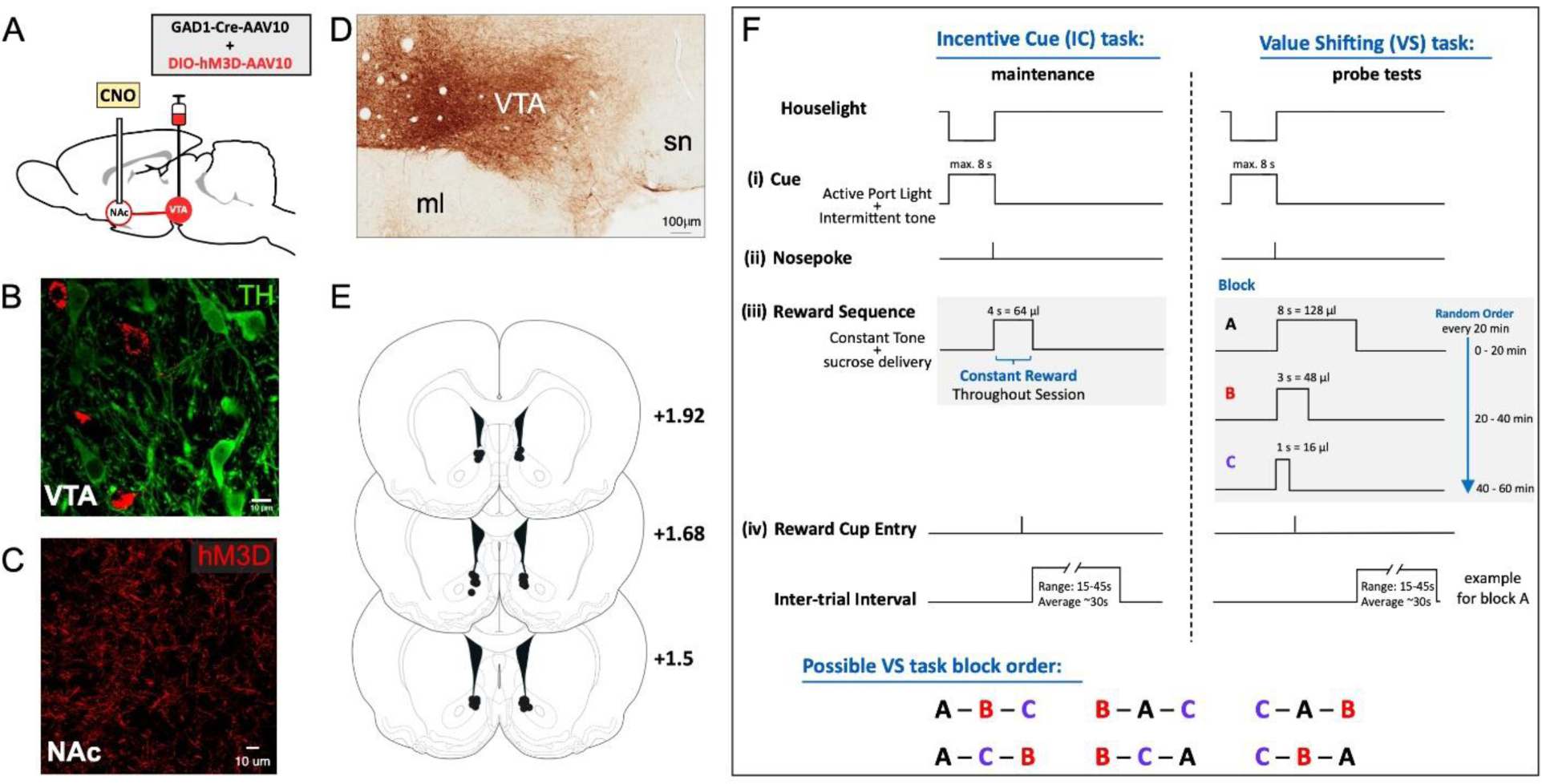
Viral targeting of activating DREADDs to VTA GABA neurons in the incentive cue and value shifting tasks. **(A)** Diagram illustrating viral targeting of VTA GABA neurons and localized activation of projections in the nucleus accumbens, **(B)** VTA neuron bodies (TH in green, hM3D in red) and (**C**) Expression of hM3D mCherry DREADD in NAc terminals. **(D)** Extent of virus expression within the VTA. **(E)** Location of the tip of the injection cannula in rats used for the VS experiments, for clarity both hM3D and tdTomato rats are shown together. **(F)** Schematics depicting the events in the incentive cue task the rats were trained and maintained on (left panel), the Value Shifting probe test (right panel), and the potential order of reward blocks in any given probe session (bottom panel). ml, medial mammillary nucleus, SN, substantia nigra, AAV10, AAV2/10.

### Behavioral Apparatus

#### Operant chambers

Med-Associates operant chambers (Georgia, VT) in sound-attenuating cubicles were equipped with two illuminated nosepoke ports on either side of a liquid receptacle (i.e. reward cup) equipped with an infrared entry detector. The left nosepoke port was designated the active nosepoke, while the right nosepoke port was designated as the inactive nosepoke. A white houselight and tone generator speaker were located on the opposite wall. Each chamber was also equipped with a pump to deliver 10% sucrose solution to the receptacle. During operant training, a 10 ml syringe was used to deliver ∼15 µl/s 10% sucrose solution, while a 30 ml syringe was used during free drinking sessions to deliver ∼45 µl/s 10% sucrose solution.

### Behavioral Training

#### Incentive cue (IC) task

Rats were trained on the Incentive Cue (IC) task, which has been modified from others and described elsewhere (18, 25-27), and in detail in the Supplemental Materials. During initial IC task training, the rats respond during a distinct 8-second audiovisual cue to obtain a ∼64 µl sucrose reward, and importantly this volume is kept constant throughout the 1 hr session (Figure 1F, left).

#### Value Shifting (VS) Task

Once rats reached criterion performance in the IC task, we probed rats with the Value Shifting (VS) task to assess the effect of altering the reward. In this task, diagrammed in Figure 1F (right), the IC predicting reward availability remained unchanged throughout all sessions. However, we divided the 1 hr session into three 20 min blocks where a successful response to the IC resulted in the delivery of one of three fixed volumes of sucrose (16 μl, 48 μl or 128 μl), which was achieved by changing the length of time the pump and associated CS+ was on (1s, 3s, 8s). There were ∼35 ICs in each block, and every IC in that 20 min block received the same designated volume of sucrose per successful response. The order of the three 20 min blocks, and the corresponding reward sizes, were randomized by the computer during the 1 hr session. All possible order permutations were equally represented in the VS probe tests (Figure 1F, bottom panel). Rats were maintained on the standard IC task (∼64 μl for 1 hr, Figure 1F, left) between VS task probes, with at least 2 days between probe sessions.

#### Free drinking, locomotor activity, and the Decreasing Reward IC task control

Additional behavioral controls are described in the Supplemental Materials.

### Histological verification, immunohistochemistry, and visualization

Immunohistochemical procedures are detailed in the Supplemental Materials.

## DATA ANALYSIS AND STATISTICAL ANALYSIS

For operant behavior, our primary metrics were the ***response ratio*** of active nosepoke responses to an IC (#nosepokes resulting in a reward/#ICs), the ***nosepoke latency*** (time; *T*) in response to the IC (T_nosepoke_ - T_IC_), and the ***latency to enter the reward cup*** (T_cup entry_ - T_nosepoke_) during the VS probe trials. We also examined a number of secondary metrics to assess whether the treatment caused behavioral changes in unconstrained responses. These include ***unrewarded active nosepokes, accuracy*** (#rewarded nosepokes/#total active nosepokes), ***active nosepokes per IC*** (#total active nosepokes/#IC), ***reward cup entries per reward*** (#total cup entries/#rewards earned), and ***total inactive nosepokes***. We adopted a sequential strategy of analysis for these experiments, which is described in detail in the Supplemental Materials.

## RESULTS

### VIRAL TARGETING VTA_(GABA)_ NEURONS

Our combinatorial viral system has been previously verified to drive transgene expression specifically in VTA_(GABA)_ neurons using electrophysiology and immunohistochemistry (IHC) (18). Briefly, when hM3D targeted neurons were activated by CNO, nearby spontaneously firing dopamine neurons were inhibited. Moreover, when targeted with channelrhodopsin-2 using a similar approach, optical stimulation produced a GABA specific optical inhibitory post synaptic current. Additionally, IHC for tyrosine hydroxylase (TH) established that hM3D expression occurred primarily in TH-neurons. Here we again confirm that hM3D-mCherry expression is distinct from tyrosine hydroxylase (TH), a marker for dopamine neurons (Figure 1B). Additionally, there is robust terminal expression of hM3D in the NAc (Figure 1C), and the spread of the hM3D-mCherry DREADD through the VTA (Figure 1D).

### ACTIVATION OF MESOACCUMBAL GABA PROJECTIONS DECREASES REWARD-SEEKING WHEN REWARD SIZE IS ABRUPTLY DECREASED

To assess how activation of mesoaccumbal GABA projections regulates reward-seeking behaviors during unpredicted changes in reward contingencies, we examined the response ratio during the VS probe task (Figure 2). When comparing the overall response ratio during each reward size across all treatments and groups, the response ratio was significantly decreased when the reward size decreased to 16 µl (Figure 2A). Three-way Mixed Effects analysis showed fixed effects of *reward* (volume) F_2,40_=23.56 p<0.0001, and *virus* (hM3D/tdTomato) F_1,40_=5.362 p=0.0258, and a *virus* x *treatment* (CNO/VEH) interaction F_1,40_=9.470 p=0.0038. Importantly, the effects on response ratio performance during the 16 µl block were specific to rats expressing hM3D and were not present with either vehicle or in DREADD-free controls. Comparing response ratios in the first and last 5 minutes of each reward bin, we found a significant decrease only in the last 5 minutes of the 16 µl reward block compared to vehicle (Figure 2B). Three-way Mixed Effects analysis of hM3D rats showed fixed effects of *reward* F_2,24_=7.016 p=0.004, *treatment* F_1,12_=15.79 p=0.0018, and *reward* x *bin* (first 5 min/last 5 min) F_2,24_=13.49 p=0.0001, *treatment* x *bin* F_1,12_=9.335 p=0.01, and *reward* x *treatment* x *bin* F_2,24_=6.081 p=0.0073 interactions. Similar comparisons in tdTomato controls showed that responding was not sensitive to CNO between the first and last 5 minutes of any reward volume. However, they remained sensitive to abrupt changes in reward volume overall (Three-way Mixed Effects analysis, fixed effects of *reward* F_2,18_=4.020 p=0.036). Moreover, tdTomato rats reduced their responding to the cue between the beginning and end of the 16 µl reward block (*reward* x *bin* interaction F_2,18_=5.222 p=0.0163).

**Figure 2.**
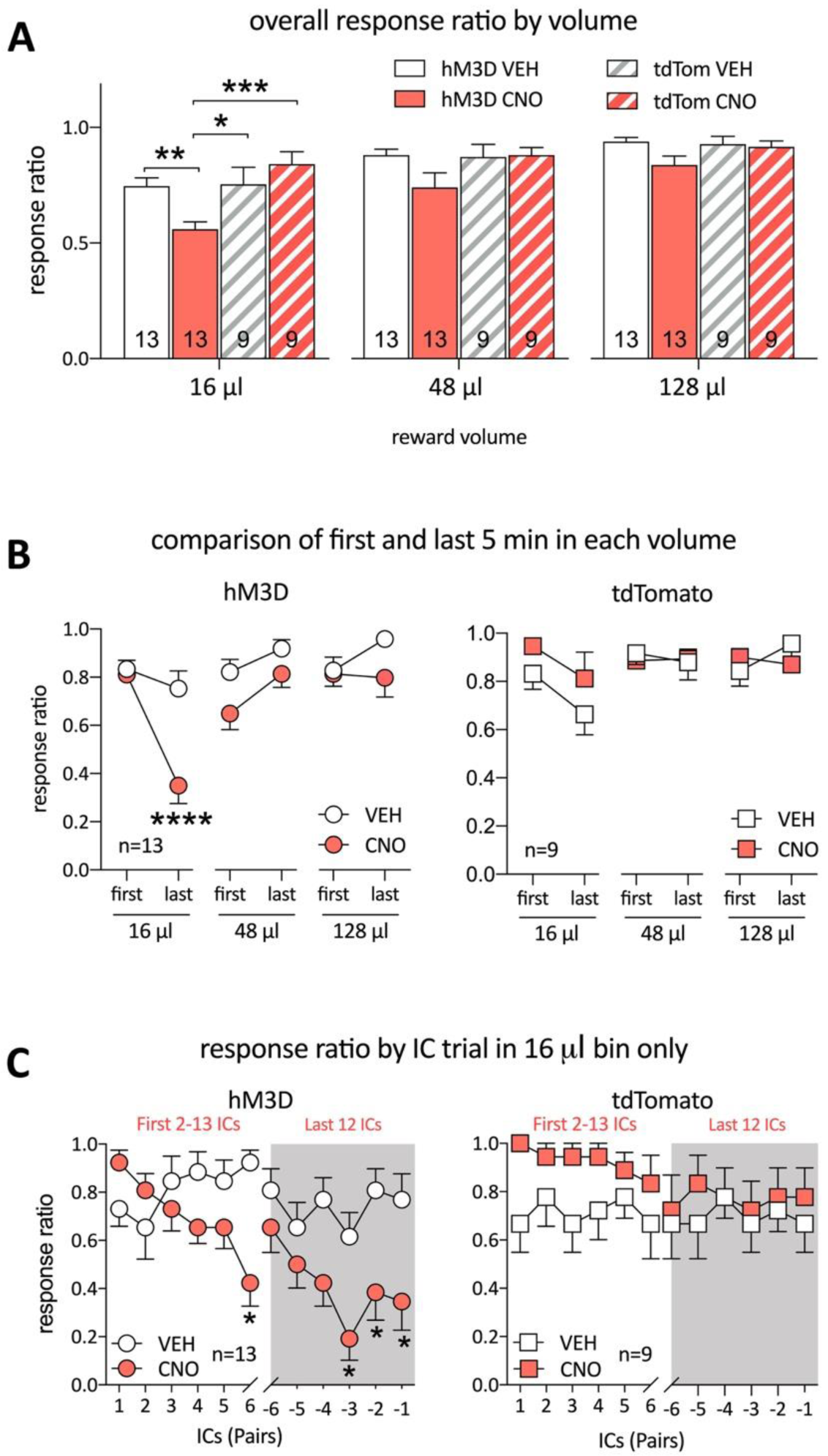
Activation of VTA(GABA) terminals in the NAc reduces responses to reward predictive cues reinforced by smaller-than-expected rewards. All data are shown as mean±SEM. (**A**) The overall response ratio significantly decreased after intra-NAc CNO in rats expressing hM3D compared to vehicle, but not in rats expressing tdTomato, when the reward size was 16 µl. **(B)** Within the first and last 5 minutes of each reward size bin, intra-NAc injections of CNO only reduced IC performance in hM3D rats at the end of the 16 µl bin.**(C)** Responses to individual successive pairs of ICs during the 16 µl bin highlights an adaptive response only in CNO treated hM3D rats after experiencing a change in reward contingency. Asterisks represent significant differences between CNO and VEH determined by a Holms-Sidak post-hoc test (*p<0.05, **p<0.01, ***p<0.005, ****p<0.001). The sample size for groups in (**A**) is found at the base of each column.

Dynamic changes in response ratio occurred on a cue by cue basis during the 16 µl reward block only after intra-accumbal CNO treatment in hM3D-expressing rats (Figure 2C, Two-way Mixed Effects analysis, fixed effects of *cue pair* F_11,132_=4.743 p<0.0001, *treatment* F_1,12_=14.88 p=0.0023, and a *cue pair* x *treatment* F_11,132_=3.325 p=0.0006 interaction). This effect was not observed in tdTomato controls.

We next examined whether activation of VTA_(GABA)_→NAc projections influences the vigor of reward-seeking in the VS probe task (Figure 3). Rats changed their latency to nosepoke to the IC relative to the reward size, and there was an overall effect of CNO treatment compared to vehicle (Figure 3A). Three-way Mixed Effects analysis showed fixed effects of *reward* F_2,40_=26.64 p<0.0001, and *treatment* F_1,20_=5.347 p=0.0315. However, there were no overall effects on *virus type*, and no significant interactions between *reward, virus type*, and *treatment*. Within the first and last 5 minutes of each reward bin, we found that nosepoke latency was significantly increased after CNO compared to vehicle only in the last 5 minutes of the 16 µl reward block (Figure 3B). In hM3D rats, there were fixed effects of *reward* F_2,24_=9.766 p=0.0008, *treatment* F_1,12_=8.911 p=0.0114, and a *reward* x *bin* F_2,24_=8.093 p=0.0021 interaction). In tdTomato controls, a similar analysis found an overall difference in nosepoke latency depending on the reward size (Three-way Mixed Effects analysis, fixed effects of *reward* F_2,16_=6.973 p=0.0066) and the nosepoke latency at the beginning and end of each reward block differed according to the reward (*reward* x *bin* interaction, F_2,16_=4.670 p=0.0253); the CNO treatment had no effect. We found no significant differences in the latency to enter the reward cup to consume reward after a successful nosepoke response to an IC, either as a function of *reward, virus type*, or *treatment* (Figure 3C). Further, there were no differences in reward cup latencies between the first and last 5 minutes of each reward bin across all of the variables tested (Figure 3D).

**Figure 3.**
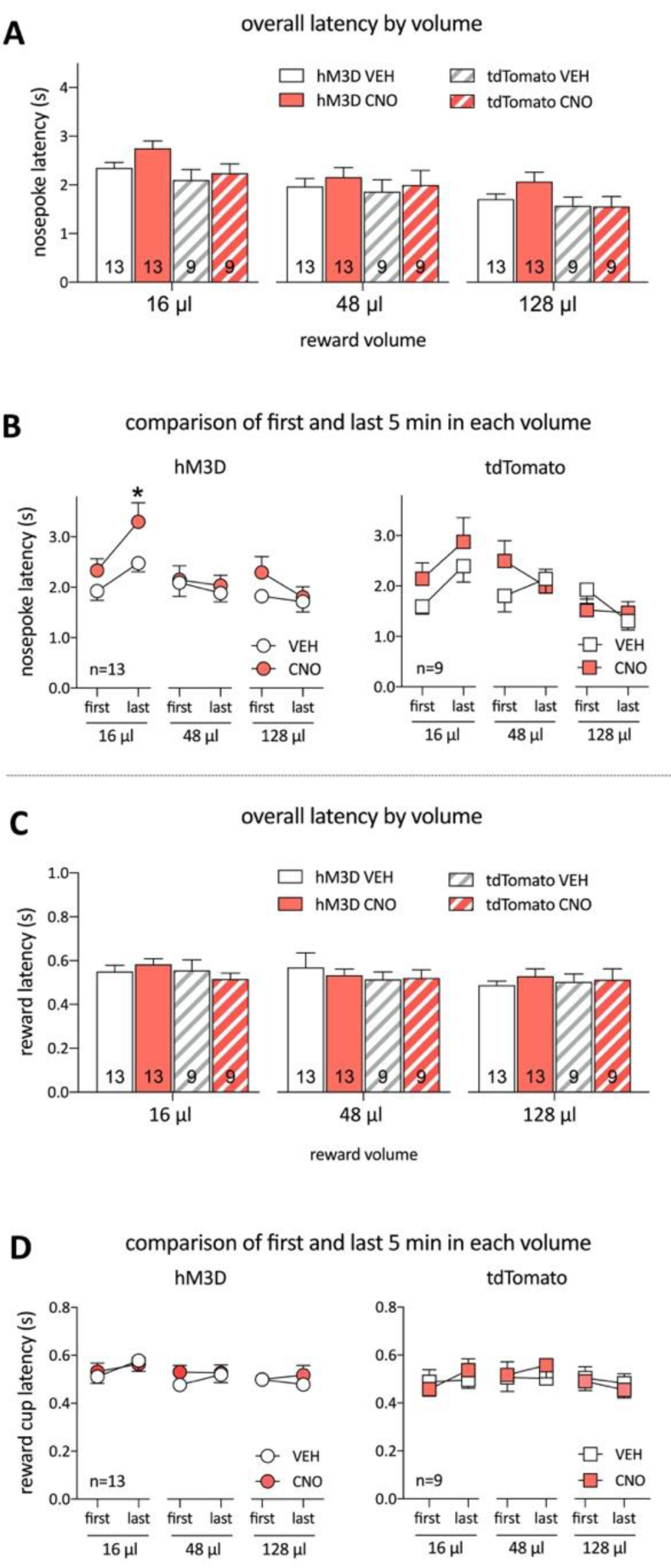
Activation of VTA(GABA) terminals in the NAc increases latency to respond to cues that predict lower than-expected-reward, but not latency to collect the reward. All data are shown as mean±SEM. (**A**) Overall latency to nosepoke increased when the reward size was reduced to 16 µl, but there were no significant differences within each reward size. **(B)** Within the first and last 5 minutes of each reward size, intra-NAc injections of CNO only increased the latency to nosepoke to the IC in hM3D rats at the end of the 16 µl bin. No effect of intra-NAc treatment was seen in latencies to enter the reward cup after responding to the IC. No within-reward size effects on latency to enter the reward cup were seen after intra-NAc treatment. Asterisks represent significant differences between CNO and VEH determined by a Holms-Sidak post-hoc test (*p<0.05)

Finally, we examined how mesoaccumbal GABA activation during the VS probe task influenced unreinforced responses, specifically unrewarded active nosepokes, accuracy, total active nosepokes per IC, reward cup entries per reward, and responses in the inactive nosepoke (Figure 4). While four of the metrics of unconstrained behavior during the VS task showed a significant effect of reward size (Three-way Mixed Effects analysis, fixed effects of *reward*, unrewarded active nosepokes: F_2,40_=7.201 p=0.0021, accuracy: F_2,40_=4.663 p=0.0151, active nosepokes per IC F_2, 40_=35.94, p< 0.0001, and reward cup entries per reward F_2, 40_=10.39 p=0.0002), demonstrating that rats were adapting their responses as reward sizes changed, there were no effects of *virus type, treatment*, or significant interactions in any group. There were no significant effects observed in inactive nosepoke responses.

**Figure 4.**
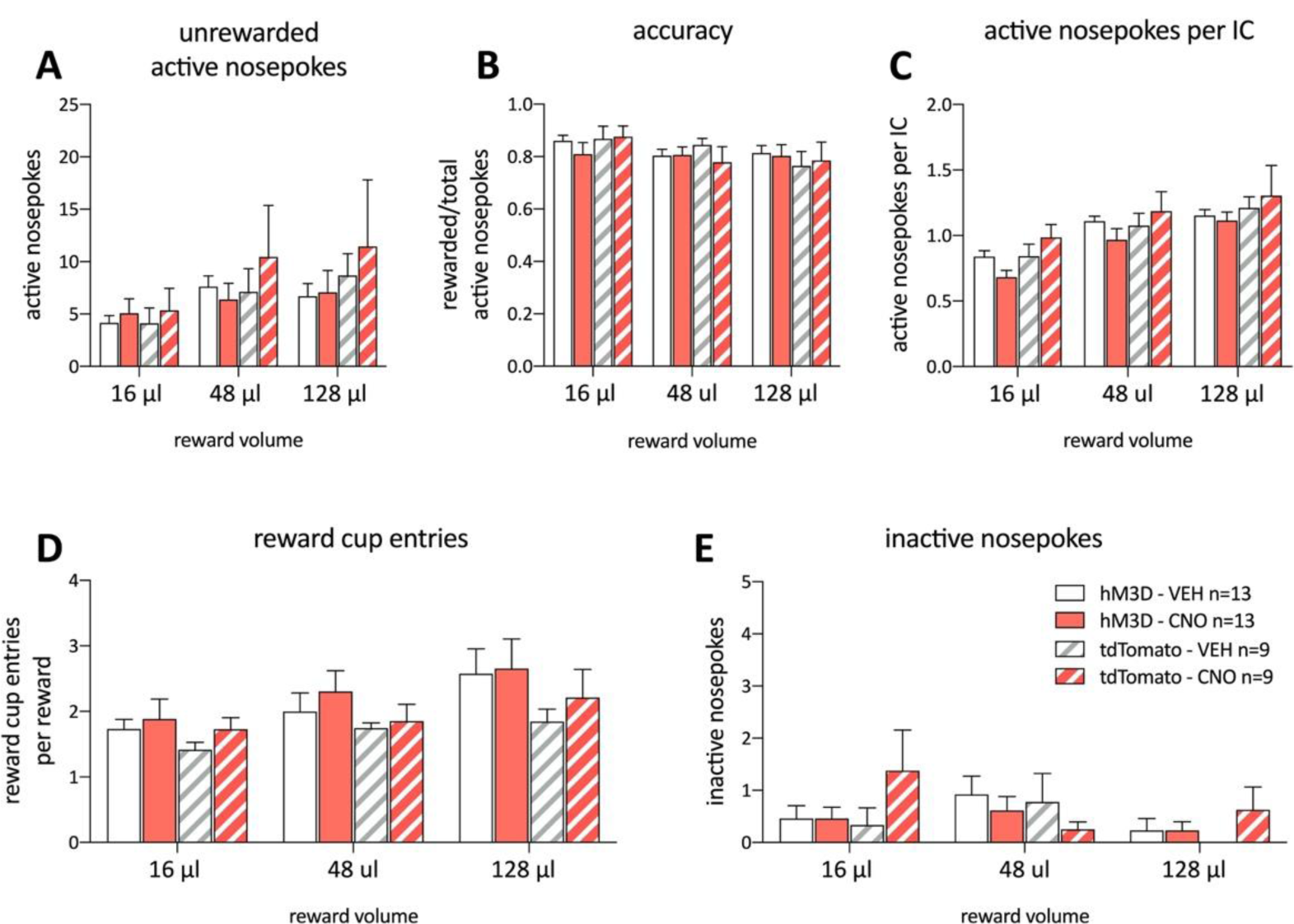
Activation of VTA(GABA) terminals in the NAc has no effect on four possible response outcomes in the VS task. All data are shown as mean±SEM. There were no significant effects of intra-NAc treatment on unrewarded active nosepokes **(A)**, accuracy (the number of rewarded nosepokes per active nosepokes) (**B**), active nosepokes per IC presented **(C)**, reward consummatory behavior as measured by reward cup entries per reward earned **(D)**, or changes in discriminating between the active and inactive nosepoke **(E)**.

### GLOBAL ACTIVATION OF VTA_(GABA)_NEURONS DECREASES REWARD-SEEKING REGARDLESS OF CHANGES IN REWARD SIZES

In contrast to targeted activation of the mesoaccumbal GABA projections, global activation of VTA_(GABA)_ neurons using systemic CNO, including local interneurons and projections to limbic and cortical regions, uniformly reduced responding to reward predictive incentive cues during the VS task (Figure 5). The response ratio was significantly decreased across all reward sizes after systemic CNO treatment (Figure5A, Three-way Mixed Effects analysis, fixed effects of *reward* F_2,40_=16.40 p<0.0001, and *virus* F_1,40_=11.14 p=0.0018, *treatment* F_1,20_=28.47 p<0.0001, and a *virus* x *treatment* interaction F_1,40_=18.36 p=0.0001). The effects of systemic CNO were limited to rats expressing hM3D. Global activation decreased the response ratio within both the first and last 5 minutes across most reward bins compared to vehicle treatment (Figure 5B, Three-way Mixed Effects analysis, fixed effects of *reward* F_2,24_=6.148 p=0.007, *bin* F_1,12_=88.41 p<0.0001, and a *reward* x *bin* F_2,24_=5.826 p=0.0087 interaction). Within the tdTomato controls we again observed a modest proportional change in response ratio with reward volume (Three-way Mixed Effects analysis, fixed effects of *reward* F_2,16_=7.827 p=0.0043), but also observed a *reward* x *treatment* x *bin* interaction (F_2,16_=6.173 p=0.0103). However, no clear trend was apparent and comparing individual responses between CNO and vehicle revealed no significant differences.

**Figure 5.**
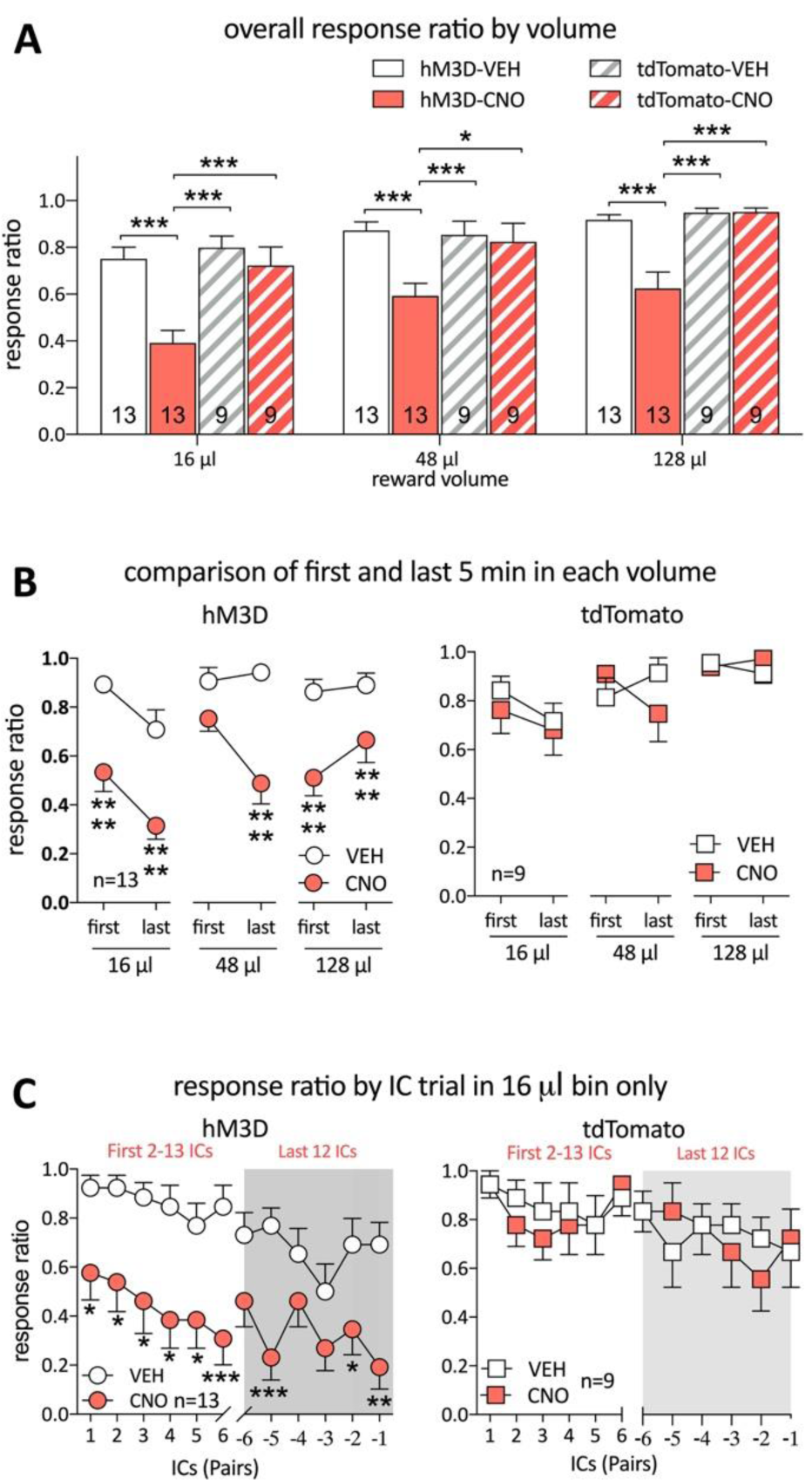
Global activation of VTA(GABA) neurons reduces responding to reward predictive cues regardless of reward size. All data are shown as mean±SEM. (**A**) Overall response ratio significantly decreased after systemic CNO in rats expressing hM3D compared to vehicle and tdTomato rats across all reward sizes. **(B)** Within the first and last 5 minutes of each reward size, systemic CNO treatment reduced IC performance in hM3D rats at both the beginning and end of the 16 µl and 128 µl reward bin. While CNO also reduced response ratios at the beginning and end of the 48 µl bin, it was only significant at the end. No differences between treatments were seen in tdTomato controls. **(C)** Responses to individual successive pairs of ICs during the 16 µl bin highlights a uniform decrease in responding to cues in CNO treated hM3D rats. Asterisks represent significant differences determined by a Holms-Sidak post-hoc test (*p<0.05, **p<0.01, ***p<0.005 ****p<0.001).

When conducting a cue by cue analysis of the response ratio performance in the 16 µl reward block, we found that global VTA_(GABA)_ activation in hM3D rats produced a uniform reduction in response ratio from the beginning of the block (Figure 5C, Two-way Mixed Effects analysis, fixed effects of *cue pair* F_11,132_=3.115 p=0.0009 and *treatment* F_1,12_=43.11 p<0.0001). In tdTomato controls, there was no effect of CNO treatment, although there was an overall modest decrease in responding on a cue by cue basis in the 16 µl reward block (Two-way Mixed Effects analysis, fixed effects of *cue pair* F_11,88_=1.932 p=0.0457).

In examining the effects of global VTA_(GABA)_ activation on other VS probe task metrics, we observed an overall increase in nosepoke latencies in all reward blocks in hM3D rats treated with systemic CNO (Figure 6A, Three-way Mixed Effects analysis, fixed effects of *reward* F_2,40_=26.78 p<0.0001, *virus* F_1,40_=8.026 p=0.0072, *treatment* F_1,20_=6.994 p=0.0155, and a *virus* x *treatment* interaction F_1,20_=7.884 p=0.0077). Subsequent analysis showed nosepoke latency was increased in hM3D, CNO treated rats compared to vehicle, between the first and last 5 minutes of each reward block (Figure 6B, fixed effects of *reward* F_2,24_=9.226 p=0.001, *treatment* F_1,12_=11.65 p=0.0051), although post-hoc tests revealed that this effect was not consistently statistically significant across all reward sizes. No effects were seen in tdTomato controls. Likewise, after global VTA_(GABA)_ activation, rats showed an increase in latency to enter the reward cup after a successful response regardless of reward size (Figure 6C, Fixed effects of *virus* F_1,40_=6.808 p=0.0127, *treatment* F_1,20_=13.39 p=0.0016, and a *virus* x *treatment* interaction F_1,40_=21.89 p<0.0001). Examining changes in the reward cup latency between the first and last 5 minutes of each reward bin showed that the latency increased regardless of whether it was at the beginning or end of the reward bin, and occurred during all reward sizes compared to vehicle (Figure 6D, Fixed effects of *treatment* F_1,12_=31.30 p=0.0001).

**Figure 6.**
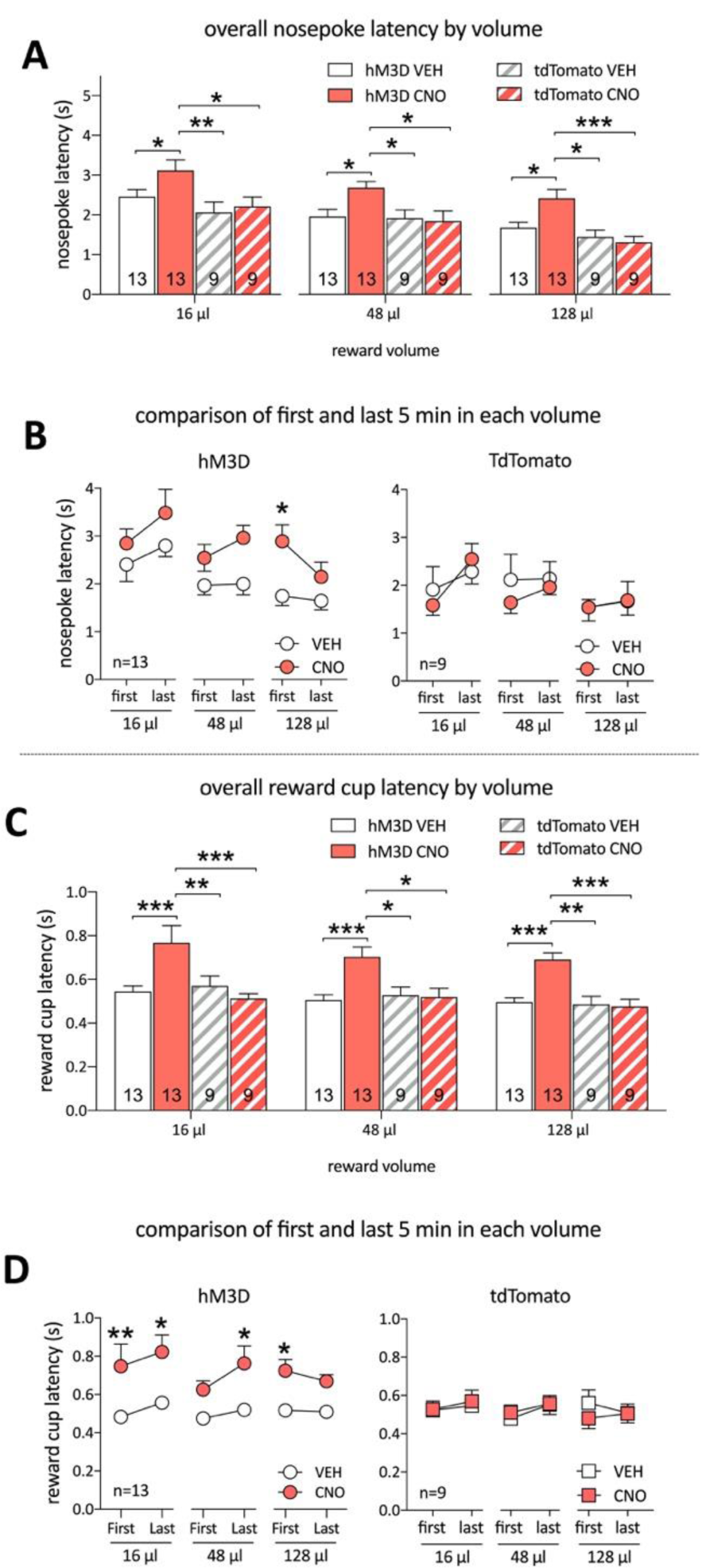
Global activation of VTA(GABA) neurons increases the latencies to respond to cues and collect the reward irrespective of reward size. All data are shown as mean±SEM. (**A**) Overall latency to nosepoke increased with all reward sizes after systemic CNO pretreatment compared to controls. **(B)** Within the first and last 5 minutes of each reward size, VTA(GABA) activation in hM3D rats increased nosepoke latencies overall in both the first and last 5 minutes of each reward bin, although most of this could be attributed to the first 5 min during the 128 µl reward block. No changes were seen in tdTomato controls. **(C)** Latencies to enter the reward cup significantly increased in hM3D rats given systemic CNO across all reward sizes compared to controls. **(D)** Latencies to enter the reward cup were significantly greater in hM3D rats during the first 5 min of the 16 and 128 µl reward blocks, and the last 5 minutes of the 48 µl bin. No within-reward block effects on latency to reward were seen in tdTomato controls. Asterisks represent significant differences determined by a Holms-Sidak post-hoc test (*p<0.05, **p<0.01, ***p<0.005).

Finally, we examined how global VTA_(GABA)_ activation during the VS probe task influenced unrewarded active nosepokes, accuracy, total active nosepokes per IC, reward cup entries per reward, and responses in the inactive nosepoke. Similar to VTA_(GABA)_→NAc projection activation, rats adapted their responses as reward sizes changed, except for responses in the inactive nosepoke (Figure 7, Three-way Mixed Effects analysis, fixed effects of *reward*, unrewarded active nosepokes, F_2,40_=6.274, p=0.0043, accuracy: F_2,40_=3.594 p=0.0367, active nosepokes per IC F_2, 40_=12.48 p< 0.0001, and reward cup entries per reward F_2, 40_=4.049 p=0.025). However, in contrast to targeted VTA_(GABA)_→NAc projection activation, global VTA_(GABA)_ activation resulted in a decrease in active nosepokes per IC after treatment across all reward blocks (Fixed effects of *virus* F_1,40_=8.002 p=0.0073, *treatment* F_1,20_=27.61, p<0.0001, and a significant *virus* x *treatment* interaction, F_1,40_ p=12.26 p=0.0012). No significant effects were observed in inactive nosepoke responses.

**Figure 7.**
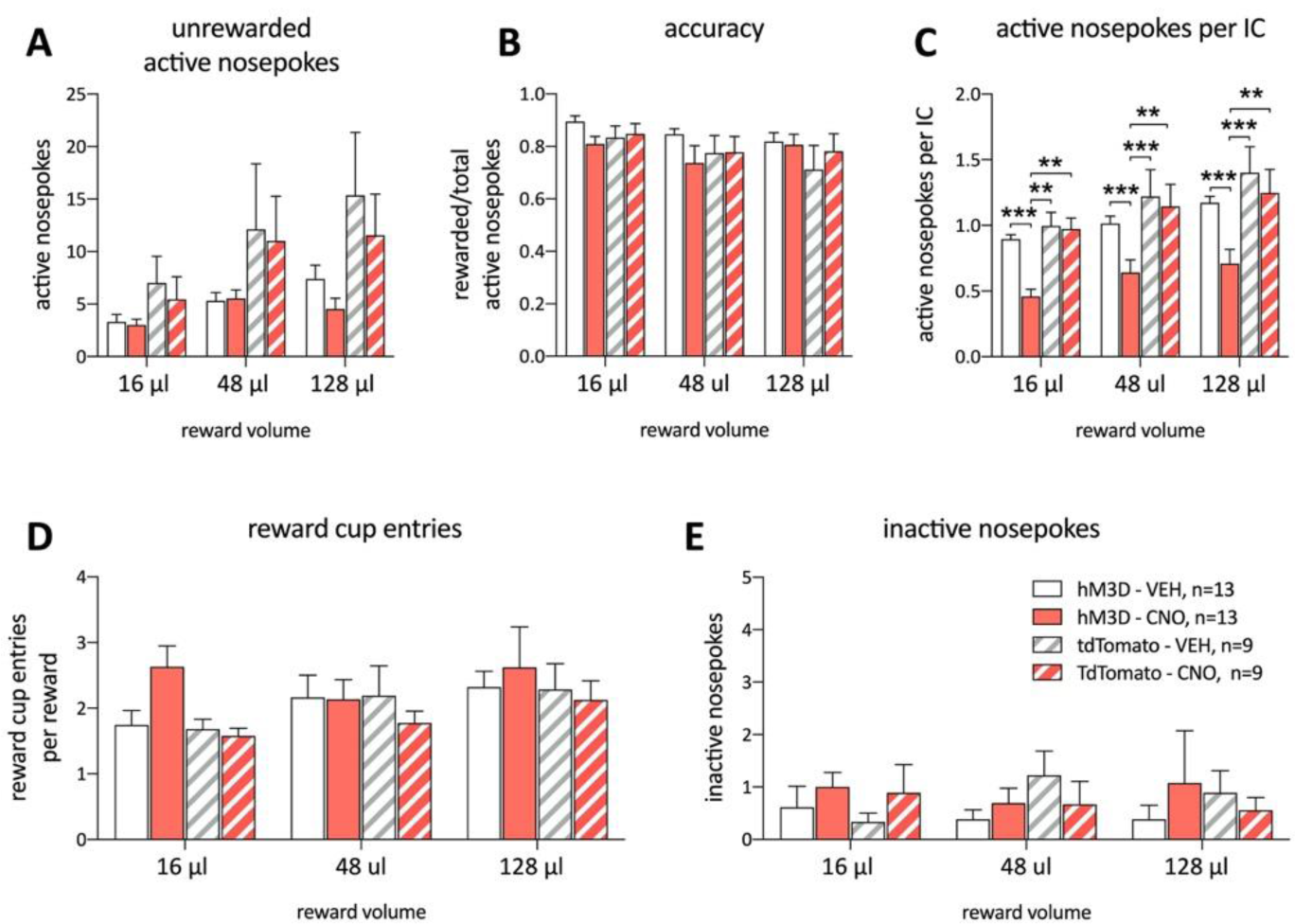
Systemic activation of VTA(GABA) neurons selectively impacts four measures of possible response outcomes to the IC in the VS task. All data are shown as mean±SEM. There were no significant effects of CNO treatment on unrewarded active nosepokes **(A)**, accuracy (the number of rewarded nosepokes per active nosepokes) (**B**), while there was a decrease in active nosepokes per IC as a result of treatment across all reward sizes compared to controls **(C)**. There were no effects of VTA-GABA activation on reward cup entries per reward earned **(D)**, or changes in the number of inactive nosepokes **(E)**. Asterisks represent significant differences determined by a Holms-Sidak post-hoc test (**p<0.01, ***p<0.005).

### MESOACCUMBAL GABA ACTIVATION DOES NOT ALTER SUCROSE CONSUMPTION OR LOCOMOTOR ACTIVITY

In a series of control experiments, we determined that mesoaccumbal VTA_(GABA)_ activation did not change the primary reinforcing effects of sucrose or baseline locomotor activity (see Supplemental Results). Further, we assessed the effect of potential conversion of CNO to clozapine (CZP) by administering an equivalent intra-NAc dose of CZP in DREADD-free controls (Supplemental Figure S1). CZP alone did not alter any of the metrics in the VS task.

### EXTENSIVE EXPERIENCE WITH REGULARLY DECREASING REWARDS NATURALLY ATTENUATES RESPONDING TO REWARD PREDICTIVE CUES

In a separate group of rats, we determined using the Decreasing Reward IC task (28), where the reward volume regularly decreases during the session, that rats naturally showed a significantly lower response ratio to 32 µl and 16 µl sucrose reward after twenty five but not one day of training (Supplemental Figure S2).

## DISCUSSION

This study was undertaken to test the hypothesis that VTA_(GABA)_ projections to the NAc specifically facilitate adapting behavioral responses in the face of changing reward contingencies. We achieved this by selectively activating mesoaccumbal GABA terminals using an activating DREADD, while changing the volume of a sucrose reward associated with a reward predictive cue. We discovered three main novel findings, first, that activation of mesoaccumbal GABA projections selectively tunes behavioral responses to predictive cues when the reward is decreased. Secondly, this attenuation in responding induced by VTA_(GABA)_→NAc projection activation occurred after experience with the altered reward, and became more pronounced over the 20 min bin. Third, the tuning was specific to responding to the IC associated with the reward, and did not impact reward consumption. These results specifically contrast our previously published data in which activation of VTA_(GABA)_→NAc projections did not affect any of these metrics when the reward volume was held constant. Additionally, we show that global activation of all VTA_(GABA)_ neurons, which affects both GABA projections to different regions and GABA interneurons, attenuated responding irrespective of experience with the new reward contingency and also attenuated reward consumption.

Reward seeking is a multifaceted process, and engages multiple reward-related systems and neural circuits (10, 29-31). Several different reward processes have been ascribed to mesolimbic circuitry, including evaluating the outcome of a goal-directed action against an established, learned value (14, 15). For example, negative reward prediction is a type of behavioral learning that occurs when an earned reward is worse than predicted (32), resulting in a subsequent decrease in seeking that reward. In our study, rats were well trained to respond to receive a fixed volume of reward (64 µl), but during VS probe tests the sucrose reward was abruptly decreased or increased from that obtained during training sessions, or in relation to larger rewards obtained in earlier reward blocks (16, 48 or 128 µl) within the session. Under these conditions, mesoaccumbal VTA_(GABA)_ activation selectively decreased the response ratio and increased the nosepoke latency to the cue predicting a much smaller volume of sucrose reward (16 µl). A central component of a learned process is that the changed reward outcome must be experienced before an alteration in reward-seeking behavior can occur. Indeed, in the VS probe rats initially responded to the cue predicting the smallest reward at levels identical to controls. Yet as they gained experience with the new reward contingency upon subsequent cue presentations, activation of VTA_(GABA)_→NAc projections facilitated a decrease in responding to the new, less-than-expected reward. In addition, the latency to respond to the cue predicting the smallest reward increased at the end of the bin, compared to vehicle controls. Together, this suggests that activation of VTA_(GABA)_→NAc projections selectively facilitated a progressive decrease in the choice to respond to the cue over time, along with a decrease in the vigor of the response (33).

Future studies will determine if these decreases facilitated by mesoaccumbal GABA activation during one session of the VS task persist when the task is repeated the next day. Also, while our data here demonstrate a decrease in responding to cues with an abrupt decrease in reward volume during the task, additional work will be needed to distinguish if this effect is due to violating established expectations of reward, or if it involves an alternative mechanism, such as potentiated reactivity to a smaller reward. It should be noted that when other rats are challenged on a Decreasing Reward IC task, where the reward always decreases reliably during the session over a similar range of volumes as the VS task, their performance during the first session is very similar to vehicle pretreated rats challenged with the VS task. This suggests that in both groups, the rats are initially unaware of the changed IC-reward contingencies and only adapt their behavior after experience. Indeed, given 25 repeated daily sessions, rats trained on the Deceasing Reward task showed similar reward-seeking behavioral adaptations as the mesoaccumbal GABA activated rats during a VS challenge. This suggests that the effects we report here are due to engaging experience-dependent processes rather an increased sensitivity to a smaller reward volume alone. It is also possible that changes in responding to the cue when rewards are unexpectedly small is secondary to altered general arousal. However, the effects of mesoaccumbal GABA activation were limited to behaviors specific to responding to the cue itself, without changes in the latency to enter the reward cup to obtain a reward, unrewarded nosepokes, accuracy, reward cup entries, inactive nosepokes, or locomotion in an open field, suggesting that altered arousal is unlikely to account for the behavior observed here.

Interestingly, a comparable increase in neither choice nor vigor of response was seen when there was an unpredicted larger-than-expected reward. It is possible that VTA_(GABA)_→NAc projections specifically contribute to forming new stimulus-outcome associations only when rewards are less than expected. However, during the development and testing of the VS task, we discovered that the task is more sensitive in detecting behavioral flexibility in the face of a smaller-than-expected reward. Therefore, we cannot preclude the possibility the rats reached a performance ceiling during this instrumental task with sucrose reinforcement, thereby masking increases in responding with larger-than-expected reward values.

To our knowledge, this is the first demonstration that VTA_(GABA)_→NAc projections can alter cue processing for rewards. We and others have generally shown that selectively activating these projections does not alter cue-induced reward seeking (17, 18). However, one study demonstrated that this pathway enhances discrimination of cues predicting an aversive stimulus (16). In that study, mice generalized freezing to two distinct tones, but optogenetic activation of VTA_(GABA)_ terminals in the NAc during conditioning resulted in the mice freezing less to the non-predictive tone, while freezing was maintained for the tone predicting the foot shock (16). In this way, the non-predictive tone was marked as less motivationally relevant. The authors determined this effect occurred through selective inhibition or “pausing” of local CINs in the NAc. Future studies examining whether our results also occur through accumbal CINs will be useful in determining if this is a common pathway for facilitating learning new cue-reinforcement contingencies regardless of valence. Additionally, future studies will be conducted to delineate the functional role of VTA_(GABA)_ terminals in the NAc core and shell subregions. It is noteworthy that neuroanatomical studies of VTA_(GABA)_ projections thus far indicate heterogenous projection patterns to these subregions (11, 34, 35) that appear distinct from dopamine.

These results substantially expand our earlier work, in which we observed no change in cue-induced responding after activating VTA_(GABA)_ projections to the NAc in the standard IC task, where reward size was kept constant (18). The VS task has different reward contingencies and is likely engaging reward-seeking processes that are distinct from those in the standard IC task. The VS probe differs notably from the IC task in that the reward sizes are randomly assigned to the same predictive cue within the session. To prevent rats from becoming over-trained on this VS task and to preserve the novelty of the changed reward contingencies, these sessions were conducted sparingly and always after several days of IC task sessions. A significant aspect of our data is that activation of VTA_(GABA)_→NAc projections altered responding in the VS challenge even though the rats were very well-trained, and continued to run in the IC task on intervening days. Thus, our subjects have a strong pre-established cue-reward expectancy due to prior training in the IC task. In contrast, after global activation of VTA_(GABA)_ neurons, we are able to show a decrease in responding, increase in nosepoke and reward latencies, and an increase in the active nosepokes per IC during the VS challenge, irrespective of the reward size. This supports our earlier findings that global VTA_(GABA)_ activation preferentially inhibits the incentive salience of the reward predictive cues (18) possibly through VTA_(GABA)_ interneuron inhibition of dopamine neurons, activity of VTA_(GABA)_ projections to other brain regions, or a combination of both effects. During the VS probe, changes in performance after global VTA_(GABA)_ activation are present at the beginning of each reward bin. This is a defining characteristic of incentive motivational processes, in that changes in reward-seeking behavior occur before experience with the reward itself (2).

Similar to our previous work, we found that the behavioral effects we report here after targeted VTA_(GABA)_→NAc activation do not result from a change in sucrose consumption under free-drinking conditions. Similarly, there was no effect of VTA_(GABA)_→NAc activation on generalized locomotion, suggesting that the decreases in responding to cues associated with a lower-than-expected reward did not result from a suppression of locomotor activity. Indeed, our data demonstrate that when the rats do respond in the other reward bins, they do so with similar speed, and are thus capable of emitting faster responses (Figure 3). In addition, since it has recently been shown that high doses of CNO are converted to CZP in vivo (36), all of our behavioral metrics for both intra-NAc CNO and systemic administration of CNO included comparisons with DREADD-free controls. We also determined that systemic CZP in DREADD-free rats did not alter the behavioral metrics of the VS probe, demonstrating that there is no confound from the potential metabolism of CNO to CZP in this study.

In summary, our data demonstrate that targeted activation of VTA_(GABA)_ projections to the NAc preferentially regulates associative learning related to new stimulus-outcomes, possibly underlying negative reward prediction processes. This specific behavioral consequence in a robust operant model is distinct from global VTA_(GABA)_ activation, which appears to preferentially regulate the attribution of incentive salience to reward predictive cues, regardless of the value of the subsequent reward. Further studies will be needed to determine the precise local circuitry within the NAc that contribute to these reward-seeking behaviors.

## Supporting information

supplemental methods and results

## Acknowledgements

This research was supported by the State University of New York BRAIN Network of Excellence Post-doctoral Fellow program and T32 AA007583 (K.T.W.) as well as The Whitehall Foundation (C.E.B.), DA043190 (C.E.B.) and AA024112 (P.J.M.). An earlier version of this manuscript has been posted on the bioRxiv preprint server (37).

