## supplemental methods and results for "Chemogenetic activation of mesoaccumbal Gamma-Aminobutyric Acid projections selectively tunes responses to predictive cues when reward value is abruptly decreased"

### SUPPLEMENTAL INFORMATION

#### *METHODS AND MATERIALS*

*Behavioral Apparatus*

Open Field chambers:

Open field chambers (43.4 x 43.4 x 30.3 cm Med-Associates, Georgia, VT) equipped with arrays of infrared sensors to detect locomotion were housed in sound attenuating cubicles illuminated by a dim white houselight.

*Incentive Cue Task Training*

Rats were initially trained on the Incentive Cue (IC) task (1-4). Briefly, rats were trained to nosepoke into the active port to receive ~64 µl 10% sucrose. In this training phase a tone and light combination (intermittent 2.9 kHz, ~80 dB tone with a 25/20 ms tone-on/off pulse, illumination of the active nosepoke port while the houselight was off) was present during the entire session, except when the rat entered the active port. This combination of stimuli comprised the IC. After a nosepoke, the syringe pump was activated for 4s, the nosepoke light was turned off, the houselight illuminated, and the intermittent tone changed to a constant tone; together these cues constituted a conditioned stimulus (CS) distinct from the IC. There was no time-out period and the session ended when the subject received 130 rewards or after 1 hr. Once rats achieved a minimum of 100 rewards for two consecutive days, rats were implanted with microinjection guide cannula and AAV virus. Rats were allowed a minimum of 3-5 days before resuming behavioral sessions. After recovery, the IC was put on a variable interval 30-s schedule, where it was presented for up to 30s and intervals between cue presentations were randomly selected from a Gaussian distribution (upper and lower limits 45s and 15s, respectively). Nosepokes into the active port during the 30s IC terminated it, resulted in reward delivery and presentation of the CS. Nosepokes into the port during the inter-trial interval (ITI), or during reward delivery, or inactive nosepoke entries, were not rewarded. Once rats responded to more than 80% of ICs during a session for two consecutive days, the IC length was shortened further to 8s, where they had the opportunity to receive ~100 rewards during the 1-hr session. Rats were run between 8-10AM, 5 days a week and trained to a performance criterion of at least 2 consecutive days of responding to ~75% of the ICs. Rats that had reached criterion performance on the IC task were limited to a maximum of 4 consecutive sessions before advancing to experimental manipulations to limit overtraining. Rats that did not achieve performance criterion in 12 consecutive days at any stage of training were removed from the study.

*Decreasing Reward Incentive Cue Task*

During training in the IC task, the sucrose reward was kept constant (64 µl) throughout the 1-hr session, while ICs in the Value Shifting (VS) task were reinforced by either 128, 48, or 16 µl of sucrose reward presented in random 20 min blocks. In the Decreasing Reward IC task, the reward volume always decreased every 15 minutes sequentially from 64, 48, 32, to 16 µl, with each block represented as previously described (5). We controlled the volume by altering the time the pump was activated, from 4s to 3s, 2s, and 1s, respectively. During all trials, the CS duration matched the pump-on time, but the length of the IC remained constant (8s). Rats (n=12) were trained on the standard IC task until acquisition (at least 2 consecutive days of responding to ~75% of the ICs), and then switched to the Decreasing Reward IC task for 25 days.

*Free drinking*

Free drinking was assessed in a subset of rats (n=8) expressing the DREADD in the $VTA_{(GABA)}$ neurons in operant chambers, as described in (1). In the free drinking task, a head entry into the reward receptacle activated the pump, delivering sucrose continuously until the rat left the reward receptacle. Chambers were checked after each session to confirm all the liquid had been consumed. Head entries in the nosepoke ports had no programmed consequences, and the houselight remained lit for the entire duration of the test. Rats were trained on this task for 3 consecutive days to acquire the task and to establish a baseline prior to commencing treatment. Rats were considered to have acquired the task once they consumed more than 0.8 ml of sucrose during the first two minutes of the free drinking session. Each session lasted a maximum of 2 hrs, and were run between 8:30AM-1:00PM. Once performance criterion was met, rats were challenged with microinfusions of CNO or an equal volume of saline vehicle as detailed above. The total volume consumed during a free-drinking session was derived from the length of time the rat remained inside the reward receptacle and the flowrate of the pump. We analyzed the first 2-min of the session as the total volume consumed in this period was equivalent to that consumed in baseline IC and VS sessions.

*Locomotor activity*

Activity was assessed in a subset of rats (n=5) expressing the DREADD in the $VTA_{(GABA)}$ neurons in open-field chambers for 1 hr. Rats were acclimatized to the chambers for two consecutive days, and pretreatment with intra-NAc CNO or saline immediately prior to the session on the third day. Locomotion was quantified as distance traveled (cm) in a 1-hr session.

*Stereotaxic Surgery*

Rats were bilaterally infused with 0.5 μl of virus combination (0.17 μl of GAD1-Cre and 0.33 μl of DIO-hM3D-mCherry AAV) into the VTA (in mm from bregma: AP -5.6, ML ±1.0, DV -8.5) over 2 min using a Nanoject III programmable injector (Drummond Scientific Company, Broomal, PA), followed by implantation with double 26-G guide cannulae (PlasticsOne, Roanoke, VA) above the NAc. Target coordinates for the cannula were (in mm from bregma): AP +1.5, ML ±1.1, DV −5.5, and were aimed towards the border of the core and shell subregions (1). The guide cannula hub was fixed to the skull with acrylic dental cement and secured with stainless steel bone screws. Obturators were inserted into the cannulae to prevent occlusions and plastic dust caps secured them into place.

*Drugs and Treatment*

To selectively activate mesoaccumbal GABA projections in the NAc, rats implanted with cannula were injected with 0.3 μl of 0.95 mM clozapine *N*-oxide (CNO, National Institutes of Drug Abuse, Baltimore, MD) dissolved in saline (6) at a rate of 0.15 μl/min, or an equal volume of vehicle. The microinjection cannulae (33-G) extended 2.0 mm beyond the ventral tip of the guide cannulae (final DV coordinate 7.5 mm below skull surface). The rats were gently restrained, the injectors inserted into the guides, and allowed to remain in place for 1 min before bilateral infusion. After the infusion, the injection cannulae were allowed to remain in place for 1 min before removal. Each rat was placed in the operant chamber for testing 15 min after the microinfusion. For global activation of $VTA_{(GABA)}$, which includes both VTA interneurons and projection neurons to the NAc, CNO was dissolved in 0.9% sterile saline and administered 0.3 mg/kg intraperitoneally (i.p.) 30 minutes prior to the start of the behavioral session. During the study, microinjection tests preceded systemic tests, and within each pair of treatments the order of CNO and saline injections were randomized. To exclude a potential confound from CNO conversion to clozapine on behavioral responses, a separate cohort of DREADD-free rats (n=6) were infused with clozapine (CZP, Enzo Life Sciences, Inc., Farmingdale, NY) dissolved with 0.9% sterile saline and 0.3 mg/kg administered i.p. 30 min prior to a VS session. These sessions were counterbalanced with systemic saline vehicle and CNO treatments.

*Statistical Analyses*

We adopted a sequential strategy for further analyses. To assess the effects of CNO or vehicle treatment on both activating DREADD (hM3D) and tdTomato (control) rats, we first directly compared the overall average response ratio and response latencies during each reward size. Subsequent tests were conducted in DREADD and DREADD-free rats in separate analyses over different time intervals. To determine if the rats were adapting their response behavior during the VS probe trials, we first compared the response ratio and response latencies during the first 5 and last 5 minutes of each reward block. We then determined the response ratio for the first twelve trials after the rats experienced the unexpected reduction in sucrose volume to the lowest volume (i.e. trials 2-13) and last twelve ICs. The first trial was excluded from each reward block as the rats were not aware of a change in sucrose volume until after they emitted their first response. Secondary metrics (unrewarded nosepokes, accuracy, active nosepokes per IC, reward cup entries per reward, and inactive nosepokes) were analyzed only with group tests without further analyses over different time intervals. Control behavioral metrics consisted of total volume consumed for free drinking sessions, distance traveled for locomotion, and response ratio during each reward volume during the Decreasing Reward IC task. All data are presented as mean ± SEM. Statistical comparisons were conducted using t-tests, one-way repeated-measures (RM) ANOVA, Two-way or Three-way mixed effects model analysis, followed by a Holms-Sidak correction for pairwise comparisons using GraphPad Prism 8. For mixed effect model analyses, we used a compound symmetry covariance matrix, fitted using Restricted Maximum Likelihood (REML). For clarity, only significant effects and interactions are reported. The level of significance was set to $\alpha=0.05$.

*Histological verification, immunohistochemistry, and visualization*

Histological verification was conducted as described elsewhere (1, 7). In brief, rats were transcardially perfused with 10% formalin and brains stored in formalin for 24-hrs at 4 °C before cryoprotecting by immersion in 30% sucrose for at least 3 days. 35 µm floating sections were obtained on a freezing microtome, separated into six series. Sections for confocal microscopy were prepared by undergoing antigen retrieval for 30 min incubation in 80 °C 10 mM sodium citrate, pH 9.0 (8), followed by washing in 0.1 M PBS containing 0.5% Triton-X (PBST) for 15 min. Sections were then pre-blocked for 2 hrs with 0.5% Normal Goat Serum. Sections were incubated for 1 hr at 23 °C, then overnight at 4 °C with primary antibody (1:1000, mouse anti-Tyrosine Hydroxylase, #22941, Immunostar). After washing three times with PBS, sections were incubated in secondary antibody (1:2000, goat anti-mouse AlexaFlor 488, A21121, Invitrogen). After washing with PBS, the steps were repeated for rabbit anti-red fluorescent protein (1:4000, #600-406-379, Rockland), and goat anti-rabbit AlexaFluor 555 (1:4000, A21429, Invitrogen). Sections were mounted, cover-slipped (Vectashield Vibrance H-1700, Vector Laboratories) and visualized with a Leica TCS SP8.

Sections for light microscopy were prepared by washing in PBST and 1% $H_2O_2$ for 30 min. Sections were then pre-blocked for 2 hrs with 0.5% Normal Donkey Serum. Sections were incubated in rabbit anti-red fluorescent protein (1:2500, #600-406-379, Rockland) overnight at 23 °C in PBST containing 0.02% sodium azide. After washing three times with PBS, sections were incubated in biotinylated secondary antibody (1:1000; 711-065-152, Jackson Immunolabs) for 2 hrs at 23 °C. Sections were washed in PBS, and incubated in avidin-biotin-horseradish peroxidase conjugate (Vector Laboratories) for 75 minutes. Sections were then washed again and incubated in a 0.06% solution of 3,3-diaminobenzidine tetrahydrochloride (DAB; Sigma, St. Louis, MO, USA) plus 0.009% $H_2O_2$. Sections were mounted, cover slipped (LiquiMount, MER77344, Mercedes Scientific), and imaged with an Apeiro ScanScope CS system.

### *SUPPLEMENTAL RESULTS*

#### *MESOACCUMBAL GABA ACTIVATION DOES NOT ALTER SUCROSE CONSUMPTION OR LOCOMOTOR ACTIVITY*

To determine if the effects we observe in the VS task may be contributable to $VTA_{(GABA)}$-induced alterations in the primary reinforcing effects of sucrose, we assessed free drinking consumption of sucrose. Under these conditions, rats will drink in 2-min a similar amount of sucrose obtained during the entire 1-hr VS task challenge at baseline or after saline. These data indicate that the consumption of sucrose remained unchanged after mesoaccumbal $VTA_{(GABA)}$ activation (Fig. S1A), and did not account for the specific changes in VS task performance within the 16 µl reward bin seen in our experiments. As well, targeted mesoaccumbal activation did not impact the total distance traveled in an open field compared to saline vehicle pretreatment (Fig. S1B), indicating that the primary results in the VS task were not the result of an inhibition in locomotor activity.

Due to recent evidence that CNO is converted to CZP in vivo (9), we also determined if any of the effects of systemic CNO on VS task performance could be attributed to activity at receptors other than hM3D. We found that an equivalent systemic dose of CZP (0.3 mg/kg, i.p.), had no impact on the four of the metrics of the VS task that showed an effect after systemic pretreatment with CNO in hM3D rats (Fig.S1C-F). In agreement with other studies on operant responding (10-13), metabolism to CZP at the dose of CNO used in this study (0.3 mg/kg, i.p.) is not attributable to the primary results in our experiments.

### *EXTENSIVE EXPERIENCE WITH REGULARLY DECREASING REWARDS NATURALLY ATTENUATES RESPONDING TO REWARD PREDICTIVE CUES*

In a separate group of rats that were initially trained on the IC task, we determined using the Decreasing Reward IC task (5), where the reward volume regularly decreases during the session, that rats did not alter their responding to the IC during the first session when reward sizes were decreased. In contrast, we found that after 25 sessions of contiguous training, rats showed a significantly lower response ratio for a 32 µl and 16 µl sucrose reward (Supplemental Figure S2, Two-way Mixed Effects analysis, fixed effects of *reward*, $F_{1,11}=17.83$ $p=0.0014$, *session #* $F_{3,33}=21.87$ $p<0.0001$, and a *reward* x *session #* interaction, $F_{3,33}=20.06$ $p<0.0001$).

### *SUPPLEMENTAL FIGURES*

Figure S1, S2

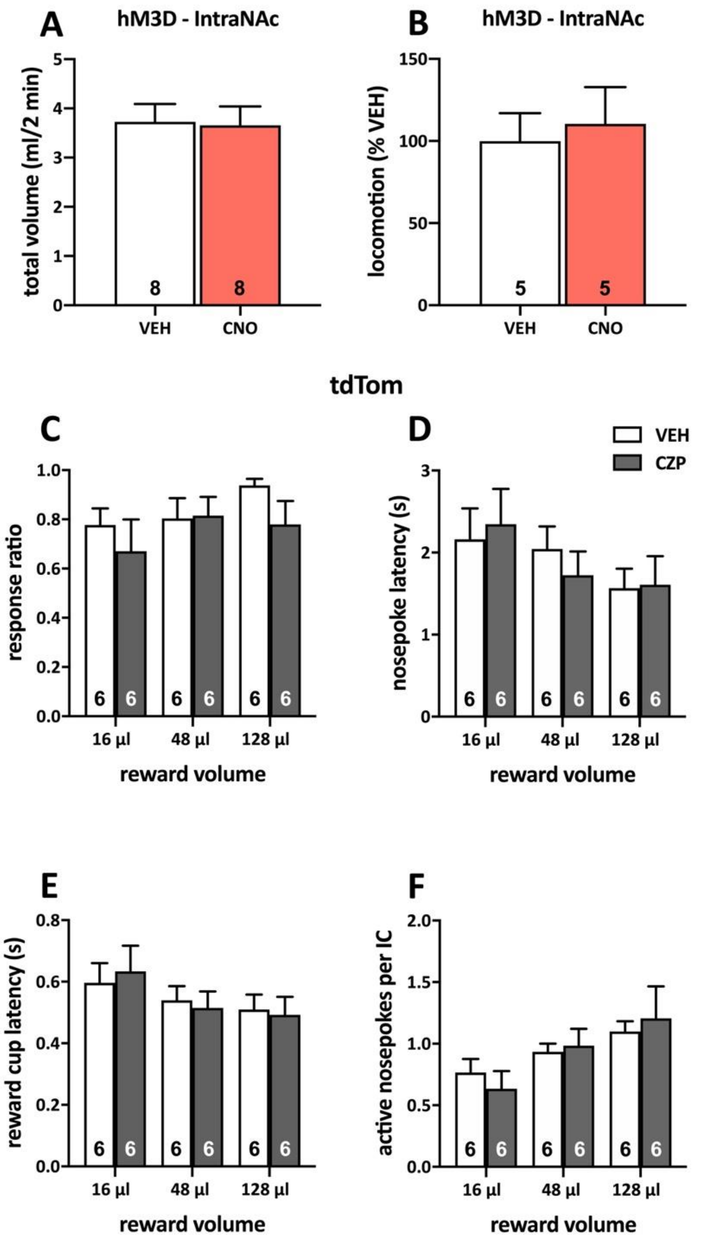


Figure S1 ***Targeted mesoaccumbal activation does not alter sucrose consumption or locomotor behavior, nor does systemic CZP impact VS task performance in DREADD-free controls.*** Data presented as mean ± SEM. **(A)** The total volume of sucrose (ml) obtained during the first 2 min of a free-drinking session produced a similar level of consumption allowed during the entire VS session. However, intra-NAc CNO pretreatment did not alter the total volume consumed during free-drinking conditions **(B).** Distance traveled (normalized as the percent of saline vehicle control) during a 1-hr open-field locomotor test also did not differ between vehicle and CNO pretreatment. Rats injected with a tdTomato virus into the VTA did not show a behavioral effect after systemic pretreatment with 0.3 mg/kg i.p. CZP in the VS task metrics of **(C)** response ratio, **(D)** nosepoke latency, **(E)** reward cup latency, or **(F)** active nosepokes per IC, four of the VS task metrics that were changed after pretreatment with CNO in hM3D rats.

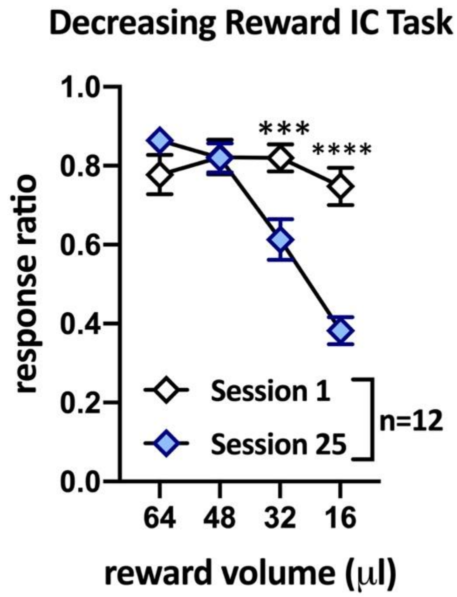


Figure S2 ***Rats naturally adapt responding to incentive cues reinforced by low reward volumes after extensive experience.*** Data presented as mean ± SEM. The response ratio during session 1 was consistent across all reward volumes. However, with time the rats adapted responding such that in session 25 the response ratio to ICs reinforced by 64 and 48 µl of sucrose was not different than in session 1, but significantly decreased when reinforced by lower volumes of 32 and 16 µl.
